# Pharmacological Macrophage Attenuation by Imatinib Reverts Hepatic Steatosis and Inflammation in Obese Mice

**DOI:** 10.1101/241224

**Authors:** Shefaa AlAsfoor, Theresa V. Rohm, Angela J. T. Bosch, Thomas Dervos, Diego Calabrese, Matthias S. Matter, Achim Weber, Claudia Cavelti-Weder

## Abstract

**Aims:** Non-alcoholic fatty liver disease (NAFLD) has become one of the most common liver diseases worldwide. As macrophages play a key role in NAFLD, therapies targeting macrophages have been postulated. Indeed, strategies depleting macrophages or blocking monocyte recruitment into the liver improve NAFLD, however, are not feasible in clinical practice. Our goal was to assess whether attenuation of macrophages can be achieved by imatinib, an anti-leukemia drug with known anti-inflammatory and anti-diabetic properties, and how this impacts NAFLD.

**Materials and Methods:** Murine macrophages were polarized *in vitro* to different activation states in the presence or absence of imatinib; mice on high fat diet orally treated with imatinib or vehicle; and human monocytes of diabetic patients and healthy controls treated with or without imatinib for translational application.

**Results:** Imatinib specifically attenuated pro-inflammatory murine macrophages *in vitro* and *in vivo*. In livers of obese mice, imatinib caused Kupffer cells to adopt an attenuated phenotype via modulation of the TNFα-pathway. This immune-modulation resulted in markedly improved hepatic steatosis along with beneficial effects on liver function, lipids and systemic inflammation. The immune-dampening effect of imatinib also prevailed in human monocytes, indicating translational applicability.

**Conclusions:** Immune-modulation of myeloid cells as exemplified by imatinib may be a novel therapeutic strategy in patients with NAFLD.

## 1. INTRODUCTION

Non-alcoholic fatty liver disease (NAFLD) is one of the most common chronic liver diseases worldwide and covers a wide disease spectrum encompassing fatty liver (steatosis), steatohepatitis (NASH) and fibrosis, even progressing to hepatocellular carcinoma.^1^ It is highly associated with metabolic disease and insulin resistance.^2^ So far, no specific treatment exists except for weight-loss, which is notoriously difficult to achieve. In recent years, evidence has accumulated that macrophages play a key role in the onset and progression of NAFLD; liver injury triggers Kupffer cell activation leading to cytokine and chemokine release, which in turn induces recruitment of inflammatory monocytes, further amplifying the disease process.^3–5^ Therefore, targeting macrophages and monocytes has been postulated as a therapeutic strategy for NAFLD.^4^ Pharmacological macrophage depletion indeed prevents the development of NAFLD in mouse models.^3,6,7^ Similarly, blocking monocyte recruitment by pharmacological or genetic ablation of different chemokine/receptor or cytokine pathways improves NAFLD characteristics (e.g. CCR2-CCL2,^5,8–10^ CCR2/5,^11,12^ CXCR3-CXCL10,^13,14^ CXCL16,^15^ IL-6,^16^ TNF-α^17^). A CCR2/5 antagonist has been tested in a clinical trial with promising results regarding fibrosis in NASH patients.^18^

Pharmacological attenuation of pro-inflammatory macrophages could be an alternative strategy to treat NAFLD. The premise of such an approach would be to specifically target pathologically activated macrophages, while not interfering with their physiological function. A potential candidate drug class with anti-inflammatory properties are PPARγ-agonists/thiazolidinediones (TZDs). TZDs are anti-diabetic drugs known to dampen pro-inflammatory macrophage activation.^19,20^ Interestingly, anti-inflammatory effects have been observed in Kupffer cells as well,^21^ potentially explaining the beneficial effects of TZDs on NAFLD.^22,23^ However, TZDs cause serious side effects such as fluid retention, congestive heart failure, weight gain and bone fractures and have therefore been largely abandoned from clinical practice. Intriguingly, the beneficial anti-diabetic/anti-inflammatory action and unwanted side effects are mechanistically distinct; while anti-diabetic/anti-inflammatory effects seem to be mediated by post-translational modification of PPARγ, unwanted side effects are a consequence of transcriptional activation of PPARγ-related genes, also termed “classical PPARγ-agonism”.^24,25^ The post-translational modification of PPARγ by TZDs involves inhibition of phosphorylation at serine273 (pS273), which can accrue with obesity and insulin resistance, and thereby restores dysregulated diabetes-related genes.^25,26^ Thus, uncoupling post-translational modification (anti-diabetic/anti-inflammatory effects) from transcriptional activation of PPARγ (side effects) could be a promising strategy for pharmacological macrophage attenuation in the setting of NAFLD.

High-throughput phosphorylation screening revealed that the tyrosine kinase inhibitor (TKI) imatinib, an established anti-leukemia drug, has specific pS273-blocking properties without classical PPARγ-agonism.^25^ Imatinib has originally been developed to target the tumor-associated fusion protein BCR-Abl in chronic myelogenous leukemia (CML). However, several other targets have been identified, pS273 being one of the most recent ones.^25,27^ Consistent with its specific pS273-blocking properties, imatinib has anti-diabetic and anti-inflammatory effects, while classical PPARγ-agonism does not occur.^25^ Anti-inflammatory effects described so far include altered immune cell polarization as shown by suppressed glycolysis in leukemia cells,^28^ adoption of an anti-inflammatory phenotype in tumor-associated macrophages,^29^ reduced inflammation in acute liver injury, TNFα-suppression in human myeloid cells,^30^ and reduced adipose tissue inflammation in an obese mouse model.^25^ Anti-diabetic effects of imatinib, observed in cancer patients and rodent diabetes models, have been postulated to arise from maintained β-cell function and reduced β-cell death.^31–35^ Based on the similarities between TZDs and imatinib regarding their anti-inflammatory effect mediated through PPARγ, an important regulator of macrophage polarization, we hypothesized that imatinib directly attenuates pro-inflammatory macrophages in disease states with pathological macrophage activation such as NAFLD. We set out a proof-of-concept study to addresses this novel therapeutic concept in a stepwise fashion: first by testing the effect of imatinib on macrophage activation *in vitro*, second on macrophages and NAFLD in metabolic disease models *in vivo*, and third on human monocytes to assess its translational application.

To specifically attenuate pathologically activated macrophages in the context of NAFLD is an intriguing concept. It implicates that a molecular switch involving the TNFα-pathway - potentially through pS273 – exists to gauge the activation state of macrophages without altering their physiological function. A more profound understanding of macrophage modulation and the molecular pathways involved will help to define its role as a therapeutic strategy in NAFLD.

## 2. MATERIAL AND METHODS

### 2.1. Animals

Male C57BL/6N mice (Charles River Laboratories) were maintained in our SPF-facility. All procedures were approved by the local Animal Care and Use Committee.

### 2.2. Murine macrophages

Peritoneal cells were harvested from 6–8-week old male C57BL/6N mice by intra-abdominal lavage, cultured overnight and enriched for macrophages by washing away non-adherent peritoneal cells. For bone marrow derived macrophages (BMDM), bone marrow cells were isolated from murine femur and tibia and differentiated by M-CSF (10 ng/mL, PeproTech) for 7–9 days. Peritoneal macrophages or BMDM were polarized to a pro- (M1; 10 ng/mL IFNγ, PeproTech, 100 ng/mL Lipopolysaccharide (LPS) E.coli0111:B4, Sigma-Aldrich) or anti-inflammatory phenotype (M2; 10 ng/mL IL-4 and IL-13, Thermo Fisher Scientific) or left non-polarized (M0) in the presence or absence of imatinib (1 μM, Novartis) for 6 hours.

### 2.3. Gene expression analysis

RNA was isolated using NucleoSpin RNA kit (Macherey Nagel) and RNeasy Plus Universal Mini kit (QIAGEN). Reverse transcription was performed with SuperScriptII Reverse Transcriptase kit (Thermo Fisher Scientific). GoTaq qPCR Master Mix (Promega) was used for real-time PCR (ViiA7, Thermo Fisher Scientific). Primer sequences (Microsynth) are listed in Suppl. Tables 1 and 2.

### 2.4. Protein expression analysis

Plasma insulin, TNF-α and IL-6 were quantified by electrochemiluminescence (MESO SECTOR S 600) using kits from MesoScale Diagnostics.

### 2.5. Acute *in vivo* inflammation model

Imatinib (100 mg/kg) or water was orally administered by gavage three times during 24h prior to a single intraperitoneal (i.p.) LPS-injection (1 mg/kg). Analysis was performed 2h post LPS and peritoneal cells and macrophages assessed by PCR.

### 2.6. Chronic *in vivo* inflammation model

Mice on high fat diet for 17–27 weeks (obesity model; HFD: 58% fat, 16.4% protein and 25.6% carbohydrate, Research Diets) and mice on HFD injected with a single dose of streptozocin at week three of HFD (diabetes model; STZ: 130 mg/kg, Sigma-Aldrich) were treated with oral imatinib (100 mg/kg) or water by gavage during the last 4–13 weeks of HFD. Insulin tolerance test (ITT) and glucose tolerance tests (GTT) were performed at 4 or 8 weeks of imatinib treatment. Macrophages were evaluated by gene and protein expression and Seahorse XF Flux analysis.

### 2.7. Seahorse XF flux analysis

Glycolysis (ECAR; extracellular acidification rate) and mitochondrial respiration (OCR, oxygen consumption rate) were measured by XF96 Seahorse Metabolic Analyzer (Seahorse Bioscience) in peritoneal macrophages *ex vivo* as previously described.^36^

### 2.8. Flow cytometry of adipose tissue macrophages

Epididymal adipose tissue was minced, digested by collagenase IV (Worthington) and DNAse I (Sigma-Aldrich) at 37°C for 25–30 min. Cells were stained with the following surface markers: DAPI, CD45 (30-F11), Siglec-F (E50-2440), F4/80 (BM8), CD11b (M1/70), CD206 (C068C2) and CD11c (N418) (all from Biolegend) and analyzed using the BD LSRII instrument (BD Biosciences) and FlowJo software (TreeStar Inc.).

### 2.9. 2-NBDG in peritoneal macrophages

Peritoneal cells were incubated at 37°C and stained with 2-NBDG (50 nM, Thermo Fisher Scientific) in FACS buffer for 30 min, washed, and stained/analyzed with the macrophage markers stated above.

### 2.10. Liver histology

Hematoxylin-eosin (H&E) and Sirius red stainings were performed according to established protocols and the NAFLD activity score (NAS)^37^ assessed in a blinded fashion. Immunohistochemistry (IHC) for F4/80, CD3, B220, and Ly-6G (antibodies in Suppl. Table 3) was performed on paraffin-embedded liver sections. For quantification of immune cells, liver sections were scanned by a Prior robot/Nikon slide scanner. Three independent visual fields were semi-automatically quantified for area fraction (F4/80) or number of cells per DAPI-positive parenchymal cells (CD3, B220, Ly-6G) using the Nikon software (NIS) tool.

### 2.11. In situ hybridization (ISH)

Mouse TNFα (VB1-10175-VT) and Emr1 (VB6-12917-VT) genes were detected in formalin fixed, paraffin embedded (FFPE), 5µm liver sections using the ViewRNA ISH system (Affymetrix) as previously described ^38^. Brightfield and fluorescent images were acquired using a laser scanning confocal microscope (LSM710, Zeiss) and Zen2 software (Zeiss) and subjected to image processing with ImageJ software.

### 2.12. Liver enzymes and lipids

Liver enzymes and blood lipids were measured in mouse plasma using a Cobas 8000 modular analyzer (Roche Diagnostics) according to the manufacturer’s protocol.

### 2.13. Human monocytes and macrophages

The human study was conducted according to the Declaration of Helsinki. Study approval was obtained from the local ethics committee. All diabetic subjects (HbA1c >6.5%) and healthy volunteers (BMI 18–25 kg/m^2^) gave written, informed consent. Detailed medical history and baseline characteristics were obtained at the day of the blood draw. Monocytes were enriched using MagniSort^®^ Human Pan-Monocyte Enrichment kit (Thermo Fisher Scientific) from peripheral blood mononuclear cells (PBMCs). Human monocytes were kept for 2h to attach and activated as above (cytokines from ImmunoTools,) in the presence or absence of imatinib (1μM) for 24h.

### 2.14. Data analysis

Data are expressed as mean±SEM. Unpaired Mann-Whitney test was used for statistical significance (GraphPad Prism). A p-value <.05 was considered as statistically significant. Correlation was calculated with Pearson’s *r*.

## 3. RESULTS

### 3.1. Imatinib attenuates pro-inflammatory macrophages *in vitro*

First, we tested whether imatinib alters macrophage activation *in vitro*. After optimization of *in vitro* readouts regarding housekeeping genes (HKGs), the optimal time point of macrophage activation and imatinib dose (Suppl. Fig. 1), we systemically exposed macrophages of distinct activation states to imatinib. In peritoneal M1-activated macrophages, imatinib attenuated multiple pro-inflammatory genes, most consistently TNFα. In accordance with this, TNFα and IL-6 protein was reduced in the supernatant of imatinib-treated macrophages (Fig. 1A). The immune-dampening effect of imatinib was more pronounced with higher pro-inflammatory M1-activation upon LPS/IFNγ as shown by iNOS expression (Fig. 1B). Accordingly, unstimulated M0 and anti-inflammatory M2 peritoneal macrophages underwent no phenotypic alteration upon imatinib treatment (Fig. 1C). To confirm its immune-dampening effect in a different macrophage population, imatinib was also tested in BMDM, where it exerted similar down-regulation of pro-inflammatory markers, although less pronounced than in peritoneal cells (Fig. 1D). Anti-inflammatory gene expression was similarly assessed and showed increased Mrc1 gene expression in M1-activated peritoneal macrophages and BMDM (Fig. 1E), enhanced Mgl1 gene expression in M2 anti-inflammatory peritoneal macrophages, and no change in unstimulated M0 peritoneal macrophages upon imatinib treatment (Fig. 1F). This demonstrates that imatinib specifically attenuates pro-inflammatory genes in M1-activated macrophages, while only increasing a few M2-markers *in vitro*.

**Fig. 1.**
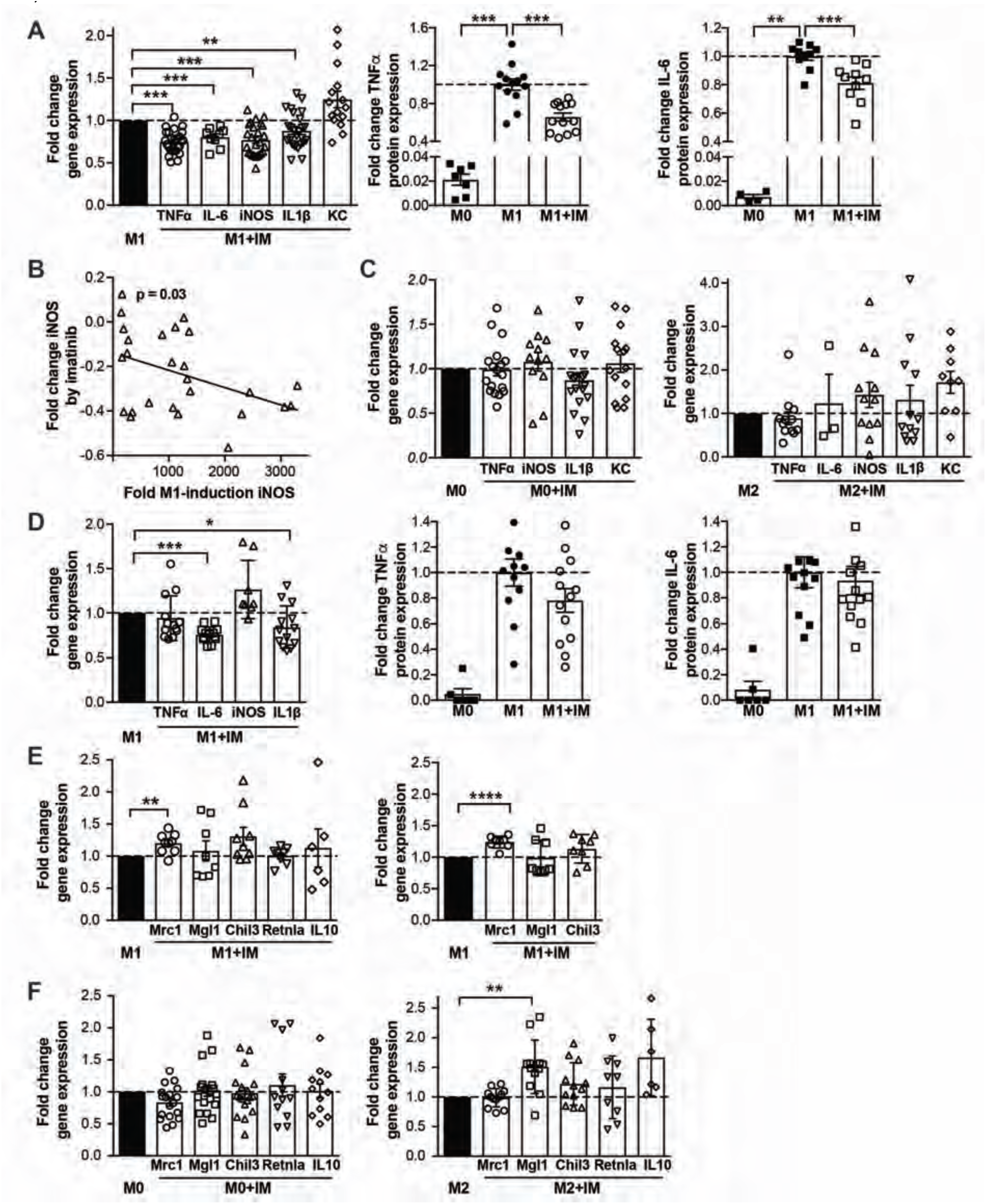
Imatinib Attenuates Pro-Inflammatory Macrophages *in Vitro*. **A:** Fold change gene expression of pro-inflammatory markers in M1-peritoneal macrophages treated with 1μM of imatinib (M1+IM, open bars) compared to non-treated M1-controls (M1, closed bar) (n = 8-26). Fold change protein expression of TNFα and IL-6 in the supernatant of unstimulated (M0), activated (M1) and concomitantly activated/imatinib-treated (M1+IM) peritoneal macrophages (n = 4-13). **B:** Correlation of M1-activation by LPS/IFNγ (expressed by iNOS-fold induction of M1-compared to M0-macrophages; x-axis) and imatinib’s immune-dampening effect (expressed by iNOS-fold change of M1+IM compared to M1-macrophages; y-axis). **C:** Fold change of pro-inflammatory gene expression in unstimulated/imatinib-treated (M0+IM, n = 12-18) and anti-inflammatory/imatinib-treated macrophages (M2+IM, n = 3-12) compared to their respective controls (M0 and M2). **D:** Fold change of pro-inflammatory gene expression in activated/imatinib-treated BMDM (M1+IM) and controls (M1) (n = 8-13). Protein expression of TNFα and IL-6 in the supernatant of unstimulated (M0), activated (M1) and concomitantly activated/imatinib-treated (M1+IM) BMDM (n = 6-13). **E:** Anti-inflammatory gene expression in M1-activated peritoneal macrophages (left) and BMDM (right; n = 6-9) treated with imatinib compared to non-treated M1-controls. **F:** Anti-inflammatory gene expression in unstimulated M0 (n = 12-18) and anti-inflammatory M2-peritoneal macrophages (n = 6-12) treated with imatinib compared to their respective controls. Data are presented as mean±SEM. *p<.05, **=p<.01, ***=p<.001.

### 3.2. Imatinib attenuates pro-inflammatory macrophages and Kupffer cells *in vivo* via modulation of the TNFα-pathway

Next, we asked whether this immune-dampening effect of imatinib on macrophages also occurs *in vivo*. To validate peritoneal macrophages as a direct macrophage readout in an acute model, we pretreated mice three times with oral imatinib followed by an i.p. LPS-injection to induce inflammation. LPS induced a highly inflammatory response with up-regulation of pro-inflammatory genes in peritoneal cells (Fig. 2A). Expression of IL-1β in peritoneal cells and TNFα in peritoneal macrophages was reduced in imatinib-pretreated mice when compared to untreated controls (Fig. 2A), thus serving as a direct readout for macrophage activation *in vivo*. This immune-dampening effect specifically concerned pro-inflammatory genes as anti-inflammatory genes were unchanged (not shown).

**Fig. 2.**
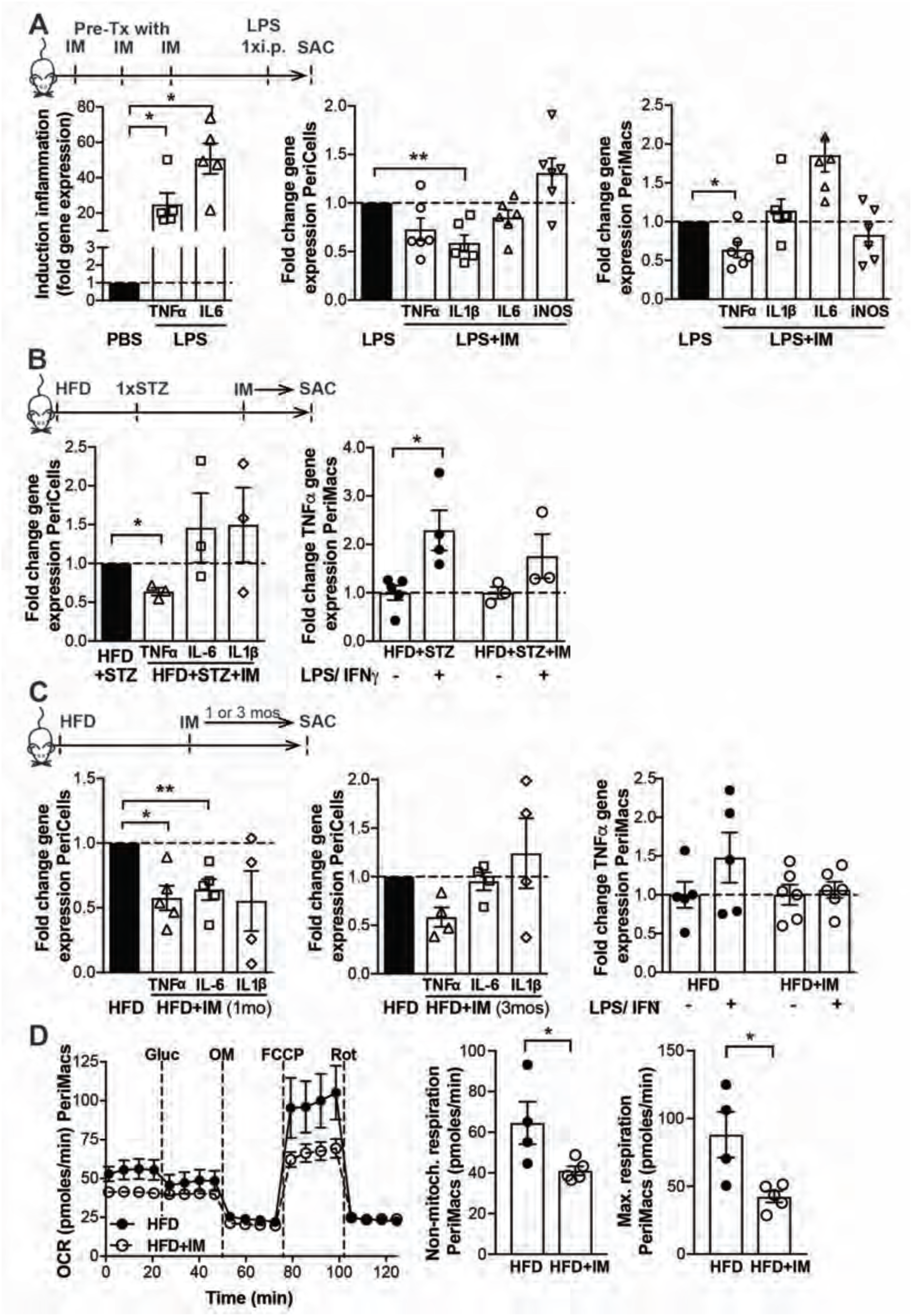
Imatinib Attenuates Pro-Inflammatory Macrophages *in Vivo*. **A:** Acute inflammation model with fold increase of pro-inflammatory gene expression in peritoneal cells from mice injected with LPS as compared to PBS-treated controls (n = 4-5).Fold change of pro-inflammatory gene expression in peritoneal cells (middle) and in peritoneal macrophages (right) of imatinib pretreated mice (LPS+IM) (n = 4-6) compared with LPS-injected controls (LPS). **B:** Fold change of pro-inflammatory gene expression in peritoneal cells (left) and in peritoneal macrophages upon additional LPS/IFNγ-stimulation (right; n = 3-6) in diabetic mice treated for one month with imatinib (HFD+STZ+IM) compared with water-treated controls (HFD+STZ). **C:** Fold change of pro-inflammatory gene expression in peritoneal cells (left: 1 month, middle: 3 months) and in peritoneal macrophages upon additional LPS/IFNγ-stimulation (right; n = 4-10) in obese mice (HFD+IM) treated with imatinib compared with water-treated controls (HFD). **D:** Seahorse flux analysis with OCR (metabolic oxidation) and calculated non-mitochondrial and maximum respiration (pmoles/min) in peritoneal macrophages of HFD-fed mice treated for 3 months with imatinib or vehicle. i.p.: intraperitoneal; PeriCells: peritoneal cells; PeriMacs: peritoneal macrophages. Data expressed as mean±SEM, *=p<.05, **=p<.01.

Next, we tested the effect of imatinib on macrophages in chronic metabolic disease. Diabetic (HFD+STZ) and obese mice (HFD) were orally treated with imatinib (IM) or vehicle and peritoneal cells harvested. TNFα gene expression was significantly reduced in peritoneal cells of diabetic mice (0.64±0.05fold) and less induced by additional LPS/IFNγ-stimulation when compared to untreated controls (1.8±0.5fold and 2.3±0.4fold, respectively, Fig. 2B). Likewise, TNFα gene expression was reduced in peritoneal cells of obese mice after one and three months of imatinib treatment (both time points 0.58±0.1fold) and less induced in peritoneal macrophages upon additional LPS/IFNγ-stimulation (1.2±0.1-fold and 2.0±0.3-fold, Fig. 2C). Similar to the acute inflammation model, anti-inflammatory genes were unaltered (Suppl. Fig. 2A,B). As an additional readout for macrophage activation, Seahorse analysis showed reduced metabolic oxidation (OCR) in peritoneal macrophages from imatinib-treated mice reaching significance for non-mitochondrial respiration and maximum respiration (Fig. 2D), while only minor effects were found for glycolysis and glucose-uptake (Suppl. Fig. 2C,D).

To test whether immune-modulation also affects Kupffer cells, which are key in the development of NAFLD, phenotypic characterization was performed. Compared to chow-fed controls, we found an increased area fraction of Kupffer cells after six months of HFD, which was reduced in mice additionally treated with imatinib in the last three months of HFD (Fig. 3A,B). In contrast, neutrophils, B- and T-lymphocytes in the liver were unaltered (Suppl. Fig. 3A,B), indicating that imatinib specifically targets macrophages. As TNFα was most affected in our previous *in vitro* and *in vivo* experiments, we probed the link between Kupffer cells and the TNFα-pathway. On a gene expression level, we found simultaneous downregulation of the macrophage marker CD68 (trend for F4/80) and TNFα in livers of imatinib-treated mice (Fig. 3C). Liver tissue hybridized with probe sets targeting Emr1 and TNFα mRNA showed co-localization of TNFα and Kupffer cells that was enhanced with HFD and reverted in imatinib-treated mice (Fig. 3D). Thus, imatinib exerts immune-dampening effects on macrophages as shown in peritoneal macrophages as a primary readout, but also causes Kupffer cells to adopt an attenuated phenotype via modulation of the TNFα-pathway.

**Fig. 3.**
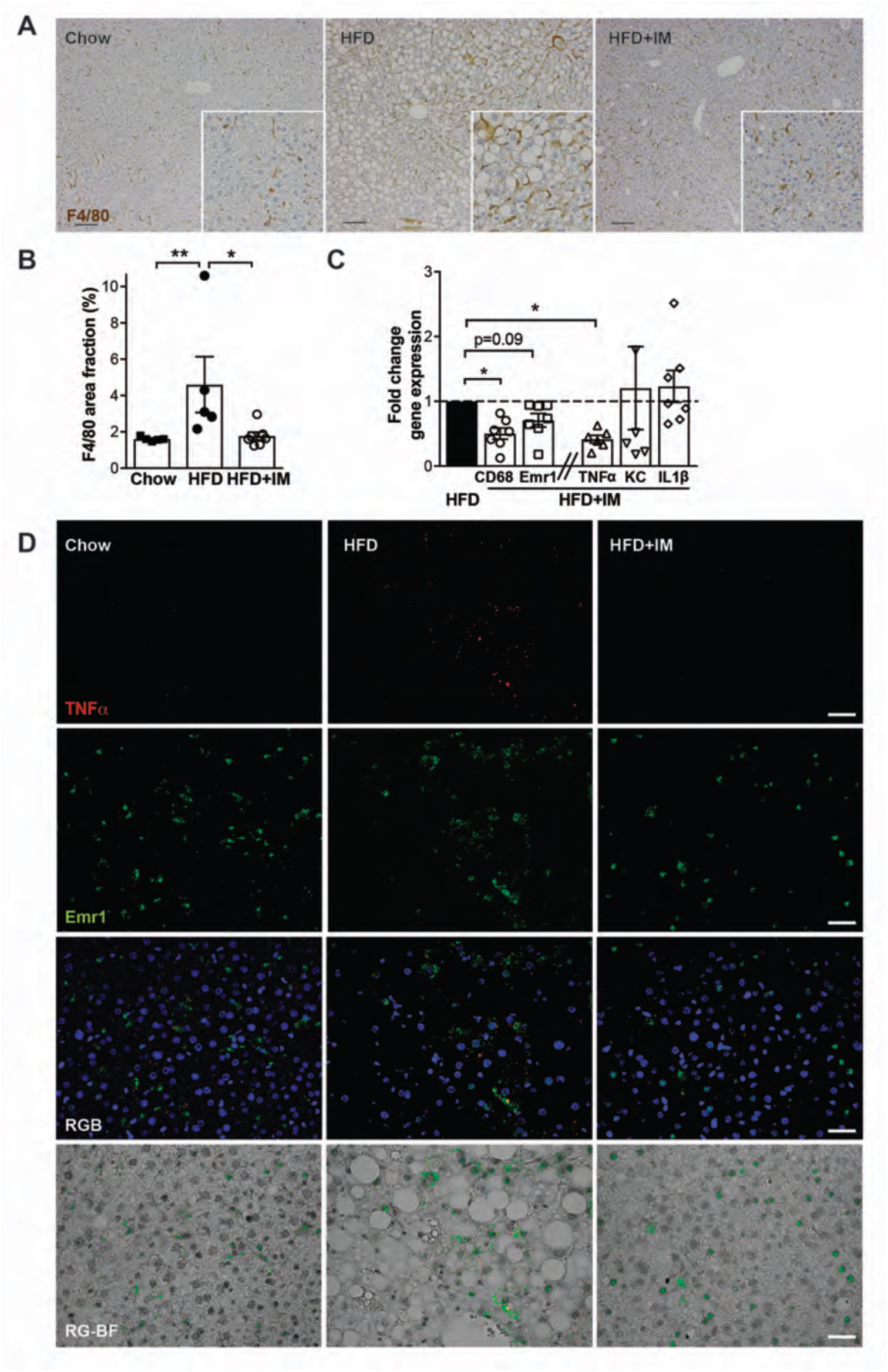
Imatinib Attenuates Kupffer Cells in NAFLD via modulation of the TNFα-pathway. **A:** Representative IHC liver sections of chow, HFD-fed and HFD+IM-treated mice stained for F4/80. **B:** Area fraction of Kupffer cells (F4/80, %) of chow, HFD and HFD+IM-treated mice. **C:** Fold change gene expression of macrophage markers (CD68, F4/80) and pro-inflammatory M1-markers (TNFα, KC, IL-1β) in HFD+IM-treated mice compared to HFD controls (n = 5-7). **D:** Mouse TNFα (red) and Emrl (green) mRNA specific in situ hybridization analysis with representative images of FFPE liver sections from chow, HFD and HFD+IM-treated mice. Red and green channel pictures were merged either with DAPI nuclear staining (RGB) or with digital bright filed (RG-BF). HFD: High fat diet; IM: imatinib. Scale bar represents 100 μm. Data expressed as mean±SEM, *=p<.05, **=p<.01.

### 3.3. Kupffer cell attenuation is associated with improved liver morphology and function

Our next question addressed whether Kupffer cell attenuation by imatinib translates to improved liver morphology and function. We found starkly increased hepatic steatosis in mice on HFD compared to chow, which was almost completely reversed after three months of imatinib treatment (Fig. 4A; Suppl. Fig. 3C). NAFLD features as quantified by the NAS-score ^37^ confirmed markedly improved liver histology in imatinib-treated mice (Fig. 4B). Fibrosis was not induced in our obesity model and therefore did not affect the NAS-score.

**Fig. 4.**
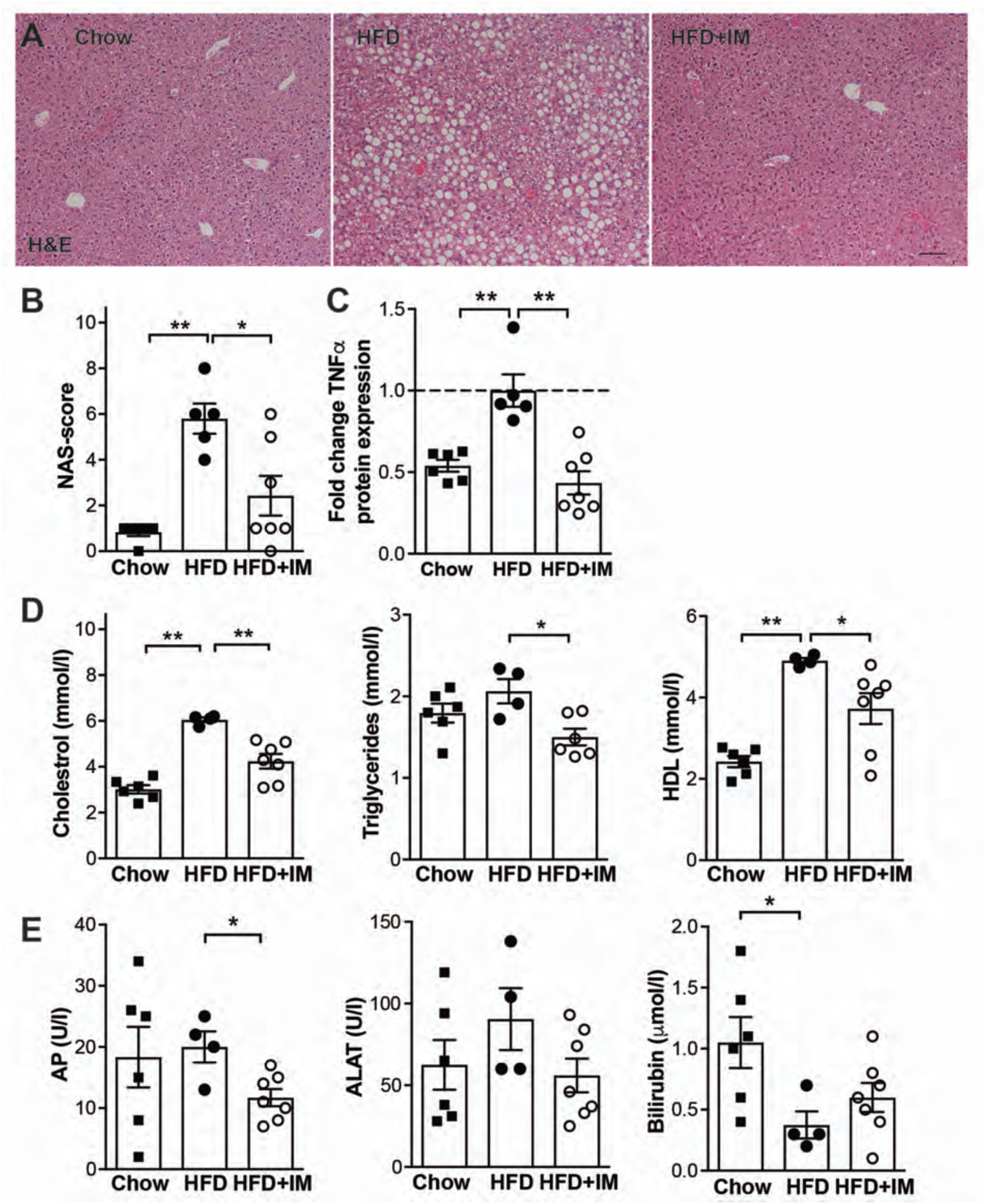
Imatinib Reverts Liver Steatosis and Improves Systemic Inflammation, Lipids and Liver Function. **A:** Representative H&E liver staining of chow, HFD-fed and HFD+IM-treated mice. **B:** Quantification of NAFLD features by the validated histological NAS-score in chow, HFD andHFD+IM-treated mice. **C:** Fold change TNFα protein expression in the blood of chow, HFD and HFD+IM-treated mice (n = 5-7). **D:** Plasma cholesterol, triglycerides and HDL in chow, HFD and HFD+IM-treated mice (n = 4-7). **E:** Liver enzymes alkaline phosphatase (AP), ALAT and bilirubin in chow, HFD and HFD+IM-treated mice (n = 4-7). HFD: High fat diet; IM: imatinib. Scale bar represents 100μm. Data expressed as mean±SEM, *=p<.05, **=p<.01.

To evaluate whether the morphological changes observed in the liver have downstream effects, we assessed blood TNFα, lipid levels and liver enzymes. TNFα was markedly decreased in imatinib-treated animals (Fig. 4C). Serum cholesterol, triglycerides and HDL - all increased with HFD - were significantly reduced in imatinib-treated mice, reflecting altered lipid metabolism (Fig. 4D). Liver enzymes showed improved liver function by significantly reduced alkaline phosphatase in imatinib-treated animals, while ALAT and bilirubin levels approached chow-fed controls in HFD-fed/imatinib-treated mice (Fig. 4E). Taken together, Kupffer cell attenuation by imatinib improves hepatic steatosis with beneficial downstream effects on systemic inflammation, lipids and liver function.

### 3.4. Imatinib reduces adipose tissue inflammation and insulin resistance

The strikingly improved NAFLD outcomes could be associated with altered macrophages in peripheral tissues and glucose homeostasis. In whole adipose tissue, macrophage and M1-markers were significantly upregulated upon HFD, which was reversed by imatinib-treatment (Fig. 5A). Flow cytometry confirmed an increased frequency of macrophages in the adipose tissue of HFD-fed animals, which was diminished upon imatinib-treatment (Suppl. Fig. 4A). Absolute numbers of adipose tissue subpopulations, however, were unchanged (Suppl. Fig. 4B-D). The frequency of eosinophils in adipose tissue was reduced in HFD-fed controls, yet an upward trend in imatinib-treated mice was found (Suppl. Fig. 4A).

**Fig. 5.**
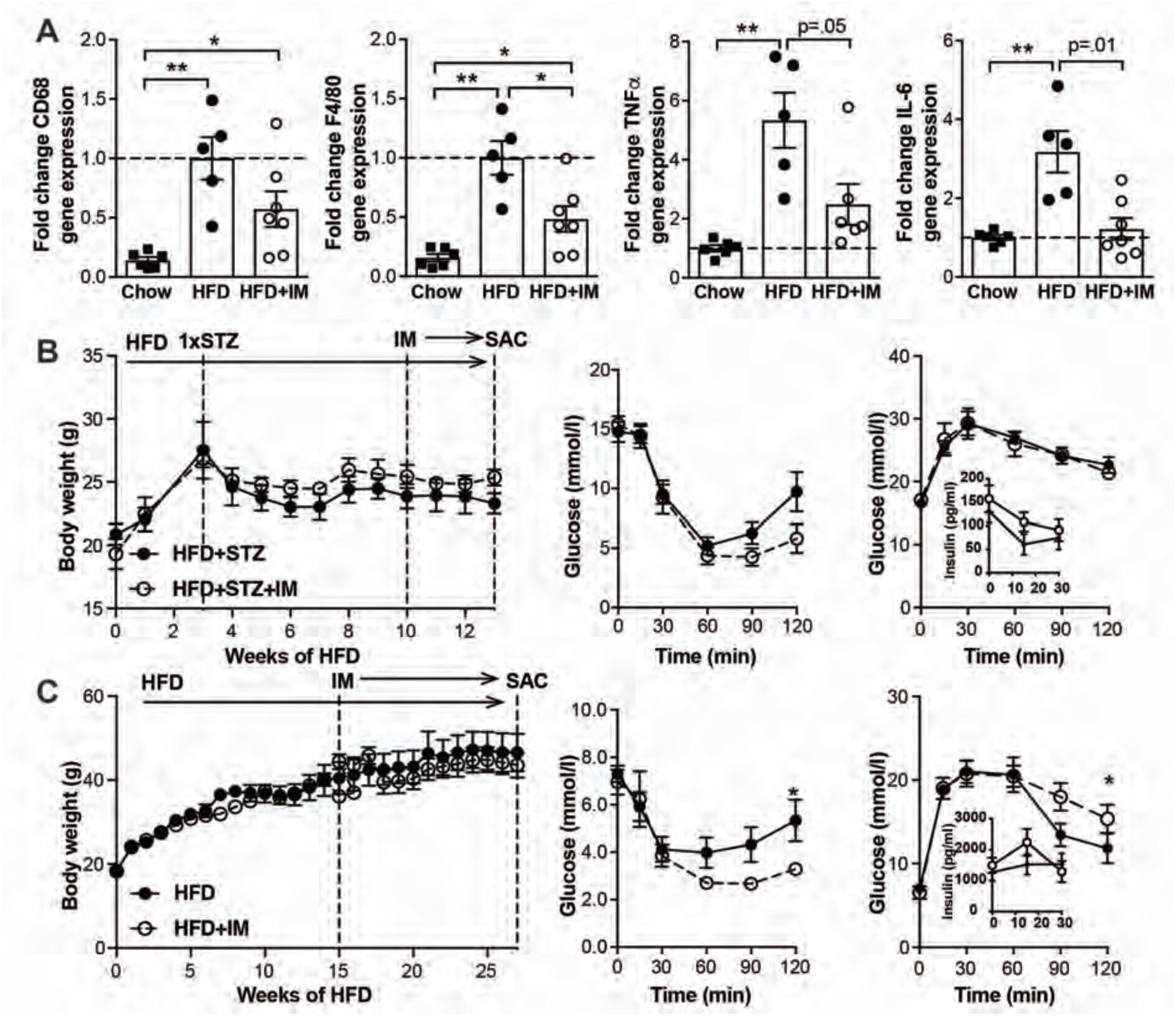
Imatinib Reduces Adipose Tissue Inflammation and Insulin Resistance. **A:** Fold change gene expression of macrophage markers (CD68, F4/80) and pro-inflammatory M1-markers (TNFα, IL-6) in whole adipose tissue of chow, HFD and HFD+IM-treated animals (n = 5-7). **B:** Body weight (left), insulin sensitivity (middle) and glucose tolerance (right; inset: insulin during GTT) in diabetic mice treated for one month with imatinib (HFD+STZ+IM) and controls (HFD+STZ; n = 4-5). **C:** Body weight (left), insulin sensitivity (middle) and glucose tolerance (right; inset: insulin during GTT) in obese mice treated for three months with imatinib (HFD+IM) and controls (HFD; n = 5-6). ATM: adipose tissue macrophages; HFD: High fat diet, IM: imatinib. Data expressed as mean±SEM, *=p<.05, **=p<.01.

**Fig. 6.**
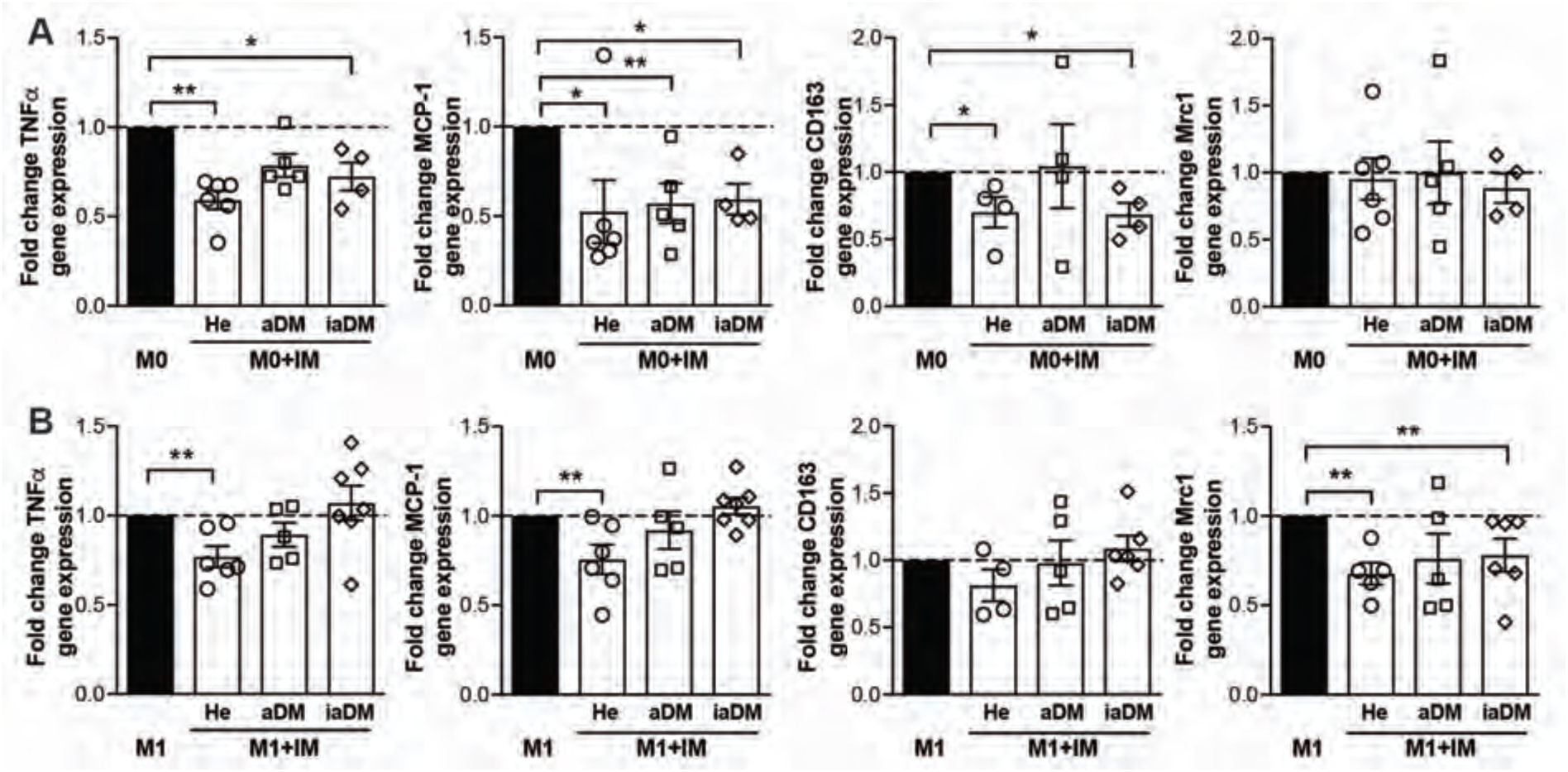
Imatinib Attenuates Human Monocytes, but Hyperglycemia Alters their Responsiveness in the Activated State. **A:** Fold change of gene expression of TNFα, MCP-1, CD163 and Mrc1 in unstimulated M0-monocytes treated with 1 μM of imatinib (M0+IM, open bars) compared to non-treated M0-monocytes (M0, closed bar) from healthy controls (He, n = 6), adequately controlled diabetics (aDM, n = 5) and inadequately controlled diabetics (iaDM, n = 5). **B:** Fold change of gene expression of TNFα, MCP-1, CD163 and Mrc1 in M1 versus M1+IM-treated monocytes from the same groups as in A. Data expressed as mean±SEM, *=p<.05, **=p<.01.

In terms of glucose metabolism, diabetic mice had slightly improved insulin sensitivity after one month of imatinib treatment when compared to vehicle-treated mice, while glucose tolerance was unaltered (Fig. 5B). Obese mice with one month of imatinib or vehicle treatment had no change in body weight or glucose and insulin tolerance. After 3 months, however, a comparable pattern as in the diabetic model was observed with improved insulin sensitivity, yet unchanged glucose tolerance (Fig. 5C). Thus, imatinib is associated with reduced adipose tissue inflammation and insulin resistance.

### 3.5. Imatinib attenuates human monocytes, but hyperglycemia alters their responsiveness

Finally, we assessed whether immune-modulation by imatinib could also be achieved in humans. As hyperglycemia is known to impact monocyte activation ^39^ and our *in vitro* data showed a differential response to imatinib depending on the activation state, we tested the effect of the drug on monocytes from subjects with markedly distinct glycemia. These included healthy controls (He), diabetics with adequate (aDM) or inadequate glycemic control (iaDM). Suppl. Table 4 shows baseline characteristics with the intended main differences between aDM and iaDM patients concerning their glycemic control (HbA1c aDM 7.0±0.5% (53mmol/mol), iaDM 12.6±0.6% (114mmol/mol)). In unstimulated monocytes, the pro-inflammatory markers TNFα, MCP-1 and CD163 were largely reduced with imatinib, while the M2-marker Mrc1 was unchanged (Fig. 5A). Interestingly, in LPS/IFNγ-activated M1-monocytes, attenuation of the pro-inflammatory markers TNFα, MCP-1 and CD163 by imatinib was gradually lost with deranged glycemic control (Fig. 5B), while the anti-inflammatory marker Mrc1 was reduced in all groups. Increasing the dose of imatinib did not rescue its immune-dampening effect in iaDM patients (Suppl. Fig. 5). In sum, unstimulated human monocytes respond to imatinib-treatment by down-regulation of pro-inflammatory markers. In activated monocytes, however, the immune-dampening effect of imatinib is lost with deranged glycemic control, suggesting altered susceptibility to the drug with hyperglycemia.

## 4. DISCUSSION

NAFLD has become one of the most common chronic liver diseases worldwide, yet no proven therapy exists. As macrophages play a crucial role in NAFLD onset and progression, we posed the question whether pharmacological macrophage attenuation improves NAFLD. As a potential immune-modulatory drug, we tested imatinib, an established anti-leukemia drug with known anti-diabetic/anti-inflammatory effects. The latter are due to imatinib’s specific pS273-blocking properties at PPARγ, which is a transcription factor regulating macrophage activation.^40^ We hypothesized that imatinib directly attenuates macrophage activation and could therefore be used in diseases with pathological macrophage activation such as NAFLD.

When first testing the notion of an immune-dampening effect of imatinib on macrophages *in vitro*, we found reduced pro-inflammatory gene and protein expression, most consistently TNFα. To validate an *in vivo* macrophage readout without an isolation or differentiation step potentially altering activation, we set up an acute inflammatory mouse model using LPS-injections. In line with our *in vitro* data, imatinib pretreatment reduced pro-inflammatory genes in peritoneal macrophages. In a similar model assessing TZDs, no immune-dampening effect had been observed when measuring serum cytokines.^41^ This indicates that direct assessment of macrophages better reflects their local activation than systemic cytokines, at least in an acute set up. The immune-dampening effect of imatinib was corroborated in chronic metabolic disease models by both reduced basal and stimulated TNFα in peritoneal macrophages. Although activated peritoneal macrophages have both enhanced glycolysis and mitochondrial oxidation,^42^ imatinib exclusively affected metabolic oxidation.

Next, we addressed whether Kupffer cells and especially the TNFα-pathway are affected by imatinib in metabolic disease and whether this translates to improved NAFLD outcomes. Thereby, quantification of Kupffer cells showed reduced area fraction by imatinib, while other immune cells were not altered in number. While gene expression showed simultaneous reductions in macrophage markers and TNFα with imatinib in whole liver tissue, co-localization of Kupffer cells and TNFα, which was dampened by imatinib, could be unequivocally demonstrated by ISH. Thus, it is conceivable, that imatinib directly interferes with the TNFα-pathway in Kupffer cells and thereby interrupts monocyte recruitment or Kupffer cell activation.

To test whether Kupffer cell attenuation affects NAFLD, HFD-induced obesity is a suitable model as it closely resembles human metabolic disease with the typical features of hepatic steatosis, systemic and adipose tissue inflammation and insulin resistance. Hepatic steatosis was markedly reduced by imatinib, which was associated with improved liver function, lipid levels and systemic and adipose tissue inflammation. Complementary to our long-term findings of morphologically improved liver steatosis, a recent study highlighted mechanistic aspects as imatinib interfered with the interaction between the MLL4 complex and PPARγ, thereby dampening steatotic target genes in short-term experiments.^43^ In terms of glycemic control, insulin resistance was decreased by imatinib, however, we found no effect on glucose tolerance. The time lag of the effect in the obesity model could be due to the time needed for PPARγ -phosphorylation to occur with HFD. Importantly, the beneficial metabolic effects were completely independent of body weight. This indicates that macrophage activation can be finely modulated via the TNFα-pathway to impact inflammation-related manifestations of metabolic disease independent of body weight.

As monocytes are increased in the circulation and liver of patients with chronic liver disease and monocyte infiltration into the liver has been recognized as a major pathogenic factor for NAFLD,^44^ we probed the effect of imatinib on human monocytes. An immune-dampening effect of imatinib on human myeloid cells has previously been demonstrated in the context of acute hepatitis,^30^ however, its effect in metabolic disease with altered glycemia has not been studied. Especially, as previous studies showed a heightened inflammatory state in human myeloid cells with hyperglycemia,^39,45^ diabetic patients could exhibit altered susceptibility to immune-modulatory drugs like imatinib. Besides the classical pro-inflammatory markers TNFα and MCP-1, we assessed CD163, whose soluble form has been associated with type 2 diabetes and insulin resistance^46^ as well as NAFLD severity.^47^ In contrast to macrophages, human monocytes exhibited responsiveness to imatinib with downregulation of TNFα, MCP-1 and CD163 also in an unstimulated activation state. This is consistent with previous studies showing that monocytes have a “pre-activated” basal condition that requires only a single stimulation with TLR ligands, while macrophages depend on a second signal to be activated.^48^ This “pre-activated” basal state of monocytes could also apply the other way around, namely that it makes them responsive to immune-dampening effects in an unstimulated state. In activated monocytes, we found a decreasing response to imatinib with deranged glycemic control; while activated monocytes of healthy controls still responded to imatinib, adequately controlled diabetics exhibited less and inadequately controlled diabetics no down-regulation of pro-inflammatory markers. Increasing doses of imatinib in inadequately controlled diabetics could not restore responsiveness to imatinib.

In conclusion, imatinib exerts an immune-dampening effect on activated macrophages and Kupffer cells via modulation of the TNFα-pathway with beneficial effects on liver morphology and function, inflammation and glycemic control. This immune-dampening effect also prevails in human monocytes. The significance of our findings lies in the scarcity of therapeutic measures available for NAFLD patients and the proof-of-principle that attenuation of macrophage activation as exemplified by imatinib improves NAFLD outcomes. As imatinib is an FDA-approved drug with a mild adverse effect profile and long-term safety record, clinical studies could be envisaged in the future. A more profound understanding of the role of macrophage attenuation in NAFLD and the molecular mechanisms involved - e.g. TNFα-modulation potentially through inhibition of PPARγ-phosphorylation - could pave the way for the development of novel therapeutic compounds in NAFLD.

## ACKNOWLEDGMENTS

We thank M. Braendle for inspiring this project, Michèle Baumann and André Fitsche for technical assistance, Alina Liedtke for help with the human study and members of the Donath and Hess laboratories for advice and feedback.

## AUTHORS CONTRIBUTIONS

Experimental design: SAA, TR, AB, TD, CCW. Experimental execution: SAA, TR, AB, TD, DC, MSM, AW, CCW. Data analyses: SAA, TR, CCW. Figure preparation, manuscript writing and editing: SAA, TR, AB, TD, DC, MSM, AW, CCW. CCW is the guarantor of this work.

## List of Abbreviations

ALAT: Alanine aminotransferase
AP: Alkaline phosphatase
ATM: Adipose tissue macrophages
BMDM: Bone marow derived macrophages
CML: Chronic myelogenous leukemia
ECAR: Extracellular acidification rate
FFPE: Formalin fixed, paraffin embedded
GTT: Glucose tolerance tests
HDL: High-density lipoprotein
HFD: High fat diet
HKG: Housekeeping genes
IHC: Immunohistochemistry
i.p.: intraperitoneal
ISH: In situ hybridization
ITT: Insulin tolerance test
LDH: Lactate dehydrogenase
LPS: Lipopolysaccharide
M-CSF: Macrophage colony-stimulating factor
NAFLD: Non-alcoholic fatty liver disease
NAS: NAFLD activity score
NASH: Nonalcoholic Steatohepatitis
OCR: Oxygen consumption rate
PBMC: Peripheral blood mononuclear cell
pS273: Phosphorylated serine273
STZ: Streptozocin
TKI: Tyrosine kinase inhibitor
TLR: Toll-like receptor
TZDs: Thiazolidinediones

